# An improved generic schema for high fidelity data linkage and sample tracing across complex multi-assay medical entomology studies

**DOI:** 10.64898/2026.05.11.724183

**Authors:** Deogratius Kavishe, Rogath Msoffe, Selemani Mmbaga, Lucia Tarimo, Fidelma Butler, Emmanuel Kaindoa, Nicodem Govella, Samson Kiware, Gerry Killeen

## Abstract

Evidence-based decision making on malaria vector control strategies increasingly rely on triangulation of data which requires informatics systems that can integrate data from complex, multi-stage studies involving mosquitoes. This manuscript describes a performance evaluation of an extended version of the generic schema underpinning the *VBDs360* platform, specifically improved to accommodate multiple distinct entomological assays spanning the field, insectary and laboratory. The utility of this extension, with respect to high-fidelity data linkage and robust sample traceability across complex entomological workflows, was evaluated through a case study conducted in southern Tanzania. Wild female mosquitoes were collected from 40 locations across a >4,000 km² area and then reared through multiple generations in an insectary before derived iso-female lineages were tested for phenotypic susceptibility to a pyrethroid insecticide. Such multi-generational lineages (*F₀* to *Fₙ* where *n* ≥ 2) were propagated to prevent non-heritable maternal effects on phenotype and produce enough progeny for standard WHO susceptibility assays. All samples were subsequently archived in a molecular laboratory, where all *F₀* specimens were tested for sibling species identity. A paper-based implementation of the extended schema enabled successful integration of 77,017 lines of data distributed across 6 different tables that spanned 3 distinct field, insectary, and laboratory workflows, implemented by three different teams working in different locations. At each step, fully independent and redundant primary and secondary keys enabled high fidelity error correction and sample tracing. Consistently perfect linkage between assay design and sample sorting data was achieved for *F_0_* wild-caught adults, with 100% of 66,108 record successfully linked between field capture and morphological categorization. This complete traceability extended to the propagation of derived *F_n_* lineages, with all 100 and 243 records from 9 adult-derived and 13 larval-derived lineages, respectively, correctly linked. Insecticide susceptibility phenotype further confirmed 100% linkage for 5,654 records between exposure history and recorded mortality outcome data in the insectary. Although such cross-cleaned linkages to sample analysis and storage data recorded by the laboratory team were not entirely perfect and could be improved, they were nevertheless of very high fidelity (97.3% (1967/2,022) for *F_0_* samples and 99.3% (437/440) for *F_n_* samples). Overall, this pilot application of the extended generic schema ensured robust data provenance and minimized transcription errors in this complex study distributed across multiple teams and locations. These findings demonstrate how this generic informatics framework may be scaled and adapted to support data integrity across diverse, large-scale, multi-team entomological research workflows.

## Introduction

Effective medical entomological research now requires not only field and laboratory expertise but also robust data management, biological sample archiving and informatics support (Alkhatib & Gaede, 2024; Kiware et al., 2016; Wang et al., 2019; Weller, 2023). Large field studies often generate diverse data, including mosquito counts and taxonomy, abundance, insecticide bioassay outcomes, genetic markers, and environmental conditions recorded during collection or experimentation. Integrating these heterogeneous datasets is challenging due to variability in data formats, evolving experimental designs and involvement of multiple teams (Borer et al., 2009). Therefore, in any well-organized study of mosquitoes, efficient and accurate informatics techniques are essential for reliable data collection and linking, as well as sample labeling and tracing (Koum et al., 2004). Indeed, robust informatics systems (Jones et al., 2006) are essential for managing complex entomological data. They enable unambiguous linkage across multiple data tables that include distinct sets of field, insectary and laboratory observations. Additionally, these systems support sample tracing as they move through the hands of several individuals or teams, each responsible for different workflow components of the study. It is also invaluable for such informatics systems (Dhawan et al., 2011) to be standardized and generalized, so they can be consistently applied to a wide diversity of studies, thus facilitating the pooling of data from multiple sources for much more powerful analyses and re-analyses.

In other domains of biology and ecology, standardized data schemas and databases have greatly enhanced data sharing and analysis (Bernstein, 2003; Jones et al., 2006; Madin et al., 2008; McIntosh et al., 2007; Page, 2008). For example, genomic and epidemiological data are curated in global repositories (e.g. GenBank, MalariaGEN, VectorBase) with controlled vocabularies (Benson et al., 2013). In the context of mosquitoes, VectorBase has long served as a genomic database for vector species, and ontological approaches have been used to design mosquito databases linking entomological and geographical data (Giraldo-Calderón et al., 2015).

Building on this foundation, a broadly generalizable informatics system was designed to address similar needs in entomological data management uses a generic schema and then initially implemented by customizing a set of standardized paper form templates to a diversity of different studies with very different goals, objectives and procedures (Kiware et al., 2016). These various early applications of the system enabled robust linking of entomological data and tracing of biological samples using two or more independent unique key identifiers per linkage. The latter were used to connect multiple separate data tables that define the experimental design and workflow used for mosquito sample collection and experimentation upon either field caught specimens or captive insectary colony stock (Kiware et al., 2016). Additionally, the system records the outcomes of sample sorting, reconstitution, and storage processes, as well as the observed attributes of the samples obtained through various laboratory tests (Kiware et al., 2016). Furthermore, a web-based platform was developed using this same generic schema. Known initially as *Mosquito Database* (*Mosquito DB)* but later renamed as *VBDs360* (https://info.vbds360.io/), this platform enables fully automated and correspondingly data linking, as well as purely electronic data entry with mobile application available in Google Play Store or via web browsers at the point of collection in the laboratory, insectary or field.

However, while this existing system is effective for relatively simple field studies, in which mosquitoes are either collected dead or killed soon after live capture, the generic schema underpinning it cannot accommodate more complex studies that continue to use live mosquitoes for further experiments. The study reported herein therefore reports on the performance of an important extension to the established *VBDs360* generic schema and derived database for entomological studies of mosquitoes that enables high fidelity tracing of mosquito data and samples over two or more discrete assays that may span multiple generations of genetically related specimens.

The challenging entomological study that motivated this extension of the *VBDs360* platform was conducted in southern Tanzania. Wild female *Anopheles arabiensis* mosquitoes were collected from 40 locations distributed across an area of more than 4,000 km^2^. Each live female was then individually propagated through several generations in a field insectary. The resulting lineages of progeny were assayed for pyrethroid insecticide susceptibility phenotype. Following the bioassay, all specimens were finally tested for species identity and archived in a central molecular laboratory. *An. arabiensis* (Diptera Culicidae), is a widely distributed malaria vector across sub-Saharan Africa that exhibits a remarkable set of evasive behaviours that allow it to minimize contact with vector control measures targeting humans indoors. Of particular importance is the ability of this species to feed on animals rather than humans (Killeen et al., 2001; Main et al., 2016; White et al., 1972), so it was hypothesized that *An. arabiensis* populations might exist in a protected conservation area that can exploit the abundant wild animal blood hosts living therein (Walsh et al., 2026). Crucially, it was hypothesized that the lack of insecticide pressure inside such large conservation areas could enable such refuge vector populations to retain greater susceptibility to insecticides than those in nearby villages, where intensive use of effectively treated bed nets over the last 18 years (Russell et al., 2010) has selected for ubiquitous resistance traits (Hancock et al., 2020).

Testing this hypothesis required tracking of individual mosquitoes from field collections of *F_0_* females, through insectary experiments upon *F_n_* progeny of live-caught field specimens that spanned multiple generations, and then molecular testing and archiving in the laboratory. To achieve this, an established informatics system based on a generic data schema and standardized collection forms (Kiware et al., 2016) was extended to enable high fidelity tracing of mosquito data and samples over two or more discrete *assays* that may span multiple generations of genetically related specimens. More broadly, this new extension of the *VBDs360* platform allows it to accommodate more complex study designs involving live mosquitoes and multiple experimental phases (e.g. breeding and successive assays) beyond the initial point of capture in the field.

## METHODS

### Generalizable revision and extension of the *VBDs360* generic schema

To enhance the generalizability of the generic schema underpinning the *VBDs360* platform, we first revised the original framework (Kiware et al., 2016) by replacing the term *Experimental Design* (ED) in the original schema with the new term *Assay Design* (AD), in which the word *Assay* is used to denote any of a number of discrete procedural units within the entomological study workflow, regardless of whether it is experimental per se or observational in nature. Each discrete assay may, for example, consist of field collections of adult or larval-stage mosquitoes or experimental assessments of insecticide susceptibility or behavioural phenotypes in the insectary or laboratory. Each discrete entomological assay comprises distinct design (AD) and sample sorting (SS) processes, just as per the original schema (Kiware et al., 2016), but allows for multiple discrete assays that are distinguished by an *Assay Number* (AYN) variable.

Second, this revised schema also introduces a distinct table for recording observed attributes of the environment the sample was collected from and/or the sample itself, with that new table linked directly to the assay design table (Figure 1). This addition of an *Environmental and/or Sample Observations* (ESO) table for each discrete assay addresses a critical gap in the earlier generic schema, where environmental data such as habitat descriptors, temperature, humidity, or other contextual variables were either embedded within assay records or entirely omitted when no appropriate linkage path was defined. The updated schema now allows these environmental and/or sample observations (ESO) to be recorded and managed independently, while nevertheless maintaining a clear 1:1 or 1:n linkage with the assay design (AD) table, respectively depending on whether one or more lines of environmental and/or sample attribute observation data were collected per line of assay design data.

**Figure 1.**
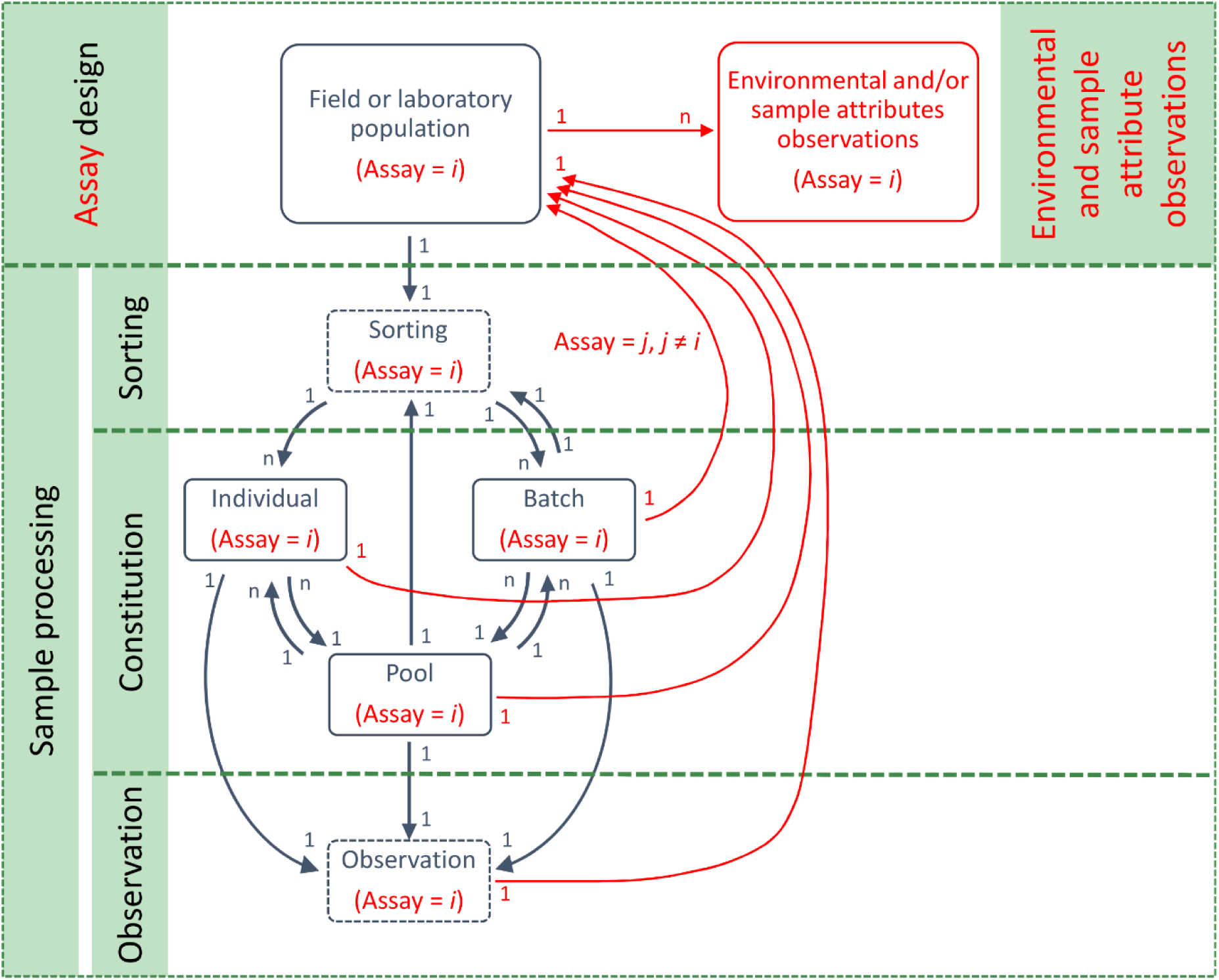
Revised schematic of the extended generic schema underpinning the *VBDs360* platform, illustrating a more generalizable framework for tracking specimens and data across multiple assays in complex entomological studies. A new Environmental and/or Sample Observations (ESO) table has been added and is linked to the *Assay Design* (AD) table through a 1: n relationship, enabling structured capture of environmental data and sample-level attributes. All new components of this extended generic schema including the indices *i* and *j* denoting the current assay (Assays=*i*), and any other distinct assay (Assay=*j*, *j*≠*i*, respectively) are distinguished by red arrows, boxes and text. The specific study that motivated this broadly generalizable extension required robust tracking of wild-caught *Anopheles arabiensis* mosquitoes all the way from field collection through to propagation and experimental phenotype assessment insectary and then molecular analysis and archiving in the laboratory (Figure 2).

This addition reflects the practical reality that many entomological studies, especially larval surveys and trap-based adult collections, often record rich environmental data that are essential for interpretation but do not otherwise fit neatly within the assay framework. Direct linking of environmental and/or sample attributes to assay design, enables more nuanced, modular integration of observational data. It also supports reuse of environmental data across multiple assays and allows greater flexibility in how observations are summarized or analysed, without forcing premature aggregation. This strategic enhancement elevates the schema’s utility beyond the context of any one study, offering a scalable model for future integration of environmental variables into complex, multi-phase entomological workflows.

### Adaptation of the generic schema to the specific motivating study

For the following motivating entomological study, the updated generic schema (Figure 1) was modified to allow entomologists to link data and samples derived from live-caught wild mosquitoes, which were used to establish derived lineages of captive-reared progeny that were then propagated through several generations to assess heritable phenotypes (Ghosh et al., 2023; Govella et al., 2023). The study-specific schema in the diagram in figure 2 shows how each of the distinct tables in which data from this study were recorded, as well as the paper forms used to collect them, are linked to each other using at least two unique identifiers per table-to-table linkage.

**Figure 2.**
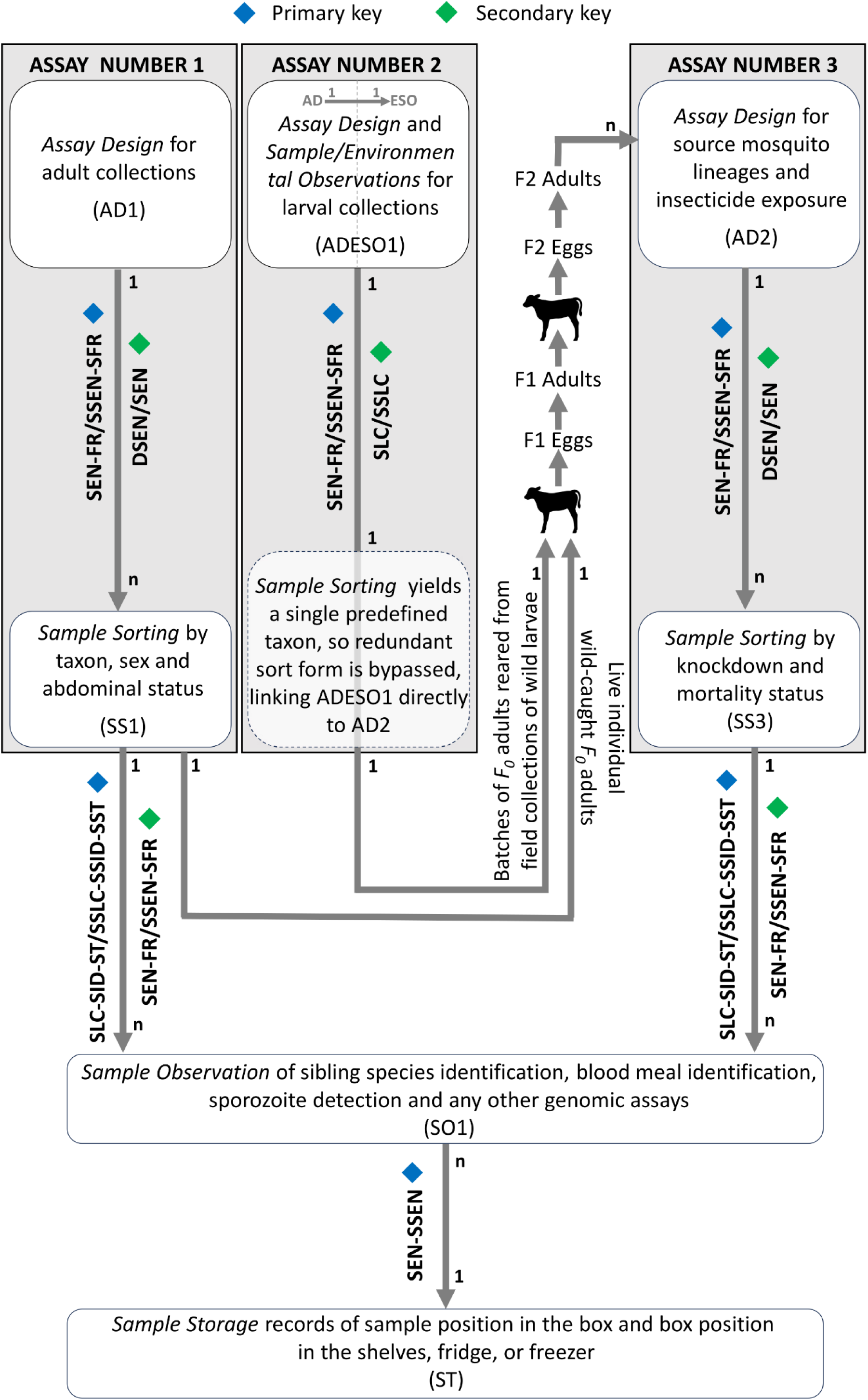
Schematic representation of the study-specific application of the extended *VBDs360* schema showing how live *Anopheles arabiensis* samples and associated data were linked across three assays: adult collections in the field (AYN=1), larval collection in the field (AYN=2) and insecticide bioassays in the insectary (AYN=3). An integrated *Assay Design and Environmental and Sample Observation* (ADESO1) form was used in Assay 2, to simplify 1:1 linkage of sampling context and habitat characteristics for collections of wild larvae. The sorting and retention of only a single, predefined larval taxon, so that the 1:1 line-by-line relationship between ADESO1 and SS data allows the redundant latter table to be bypassed, thus enabling simplified onward linkage direct to Assay 3 (AD2). All records and specimens were linked via primary and secondary keys, each comprising at least two sets of unique identifiers that were deliberately designed to be fully independent and redundant, to ensure robust cross-verification and correction. This was intended to ensure uninterrupted traceability, from initial collection of wild mosquitoes in the field all the way through to final molecular analysis (SO1) and long-term archiving (ST) in the laboratory, despite all the usual trivial human errors that occur with such paper-based systems.

Adapting the new extension of generic schema to this particular study allows three distinct entomological procedures to be distinguished by the new *assay number* variable (AYN = 1, 2 or 3). The field collections of adults in the first assay (AYN =1) were recorded in well-standardized tables and forms (AD1 linked to SS1) that were fully consistent with Figure 1 and even the preceding version of the generic schema (Kiware et al., 2016). However, the exact procedures used in the second assay (AYN = 2) to record field-based collection (AD) and sorting (SS) of larval samples, the observed environmental attributes of the aquatic habitats they were collected from (ESO), and then their subsequent propagation in the insectary allowed for some convenient bespoke simplifications of this component of the study-specific schema (Figure 2).

First, a line-by-line 1:1 linkage between the AD and ESO tables allowed all the relevant variables to be readily recorded on a single integrated *Assay Design and Environmental and/or Sample Observation* (ADESO1) form (Figure 2). Second, it was possible to entirely eliminate the need for a *Sample Sorting* (SS) form or data table of any kind, simply because the sorting process carried out in the field for this second assay was designed to yield only a single, predefined taxonomic and life stage category: Any larvae that visually appeared to resemble *An. gambiae sensu lato*, with all other mosquito larvae being discarded. Because of this 1:1 relationship with derived batches and pools of sorted mosquito larvae, it was possible for the primary and secondary keys linking the ADESO1 data onward to the next procedural stage to completely bypass the sample sorting (SS) table, which was correspondingly rendered redundant along with the corresponding forms (Figure 2). The ADESO1 table therefore links directly to the assay design table and form (AD2) for the third assay in the study design (AYN = 3), using the same primary and secondary keys to track lineage-specific progeny all the way through all the intermediary sample sorting and subsequent propagation processes that took several weeks and sometimes even months to complete in the field insectary.

### Study setting and sampling frame for the collection of wild *F_0_* generation mosquitoes

The medical entomology study that motivated the development of this extension to the *VBDs360* platform was conducted across the Ifakara-Lupiro-Mang’ula Wildlife Management Area (ILUMA WMA), some nearby villages to the west, and extensive adjacent areas of Nyerere National Park (NNP) to the east (Figure 3). Detailed descriptions of the study area and sampling procedures have been previously described in detail elsewhere (Kavishe et al., 2025A; Kavishe et al., 2025B; Walsh et al., 2026). In brief, both adult mosquitoes and their aquatic larval stages were collected at or close to each of 40 mobile camps distributed across the study area. Four rounds of such mosquito surveys were conducted between January 2022 and December 2023 (Kavishe et al., 2025A; Kavishe et al., 2025B; Walsh et al., 2026), and each was complemented by detailed surveys of all signs of activities by humans, livestock and wild animals, all of which represent potential sources of blood for mosquitoes (Duggan et al., 2024; Duggan et al., 2025; Walsh et al., 2026).

**Figure 3.**
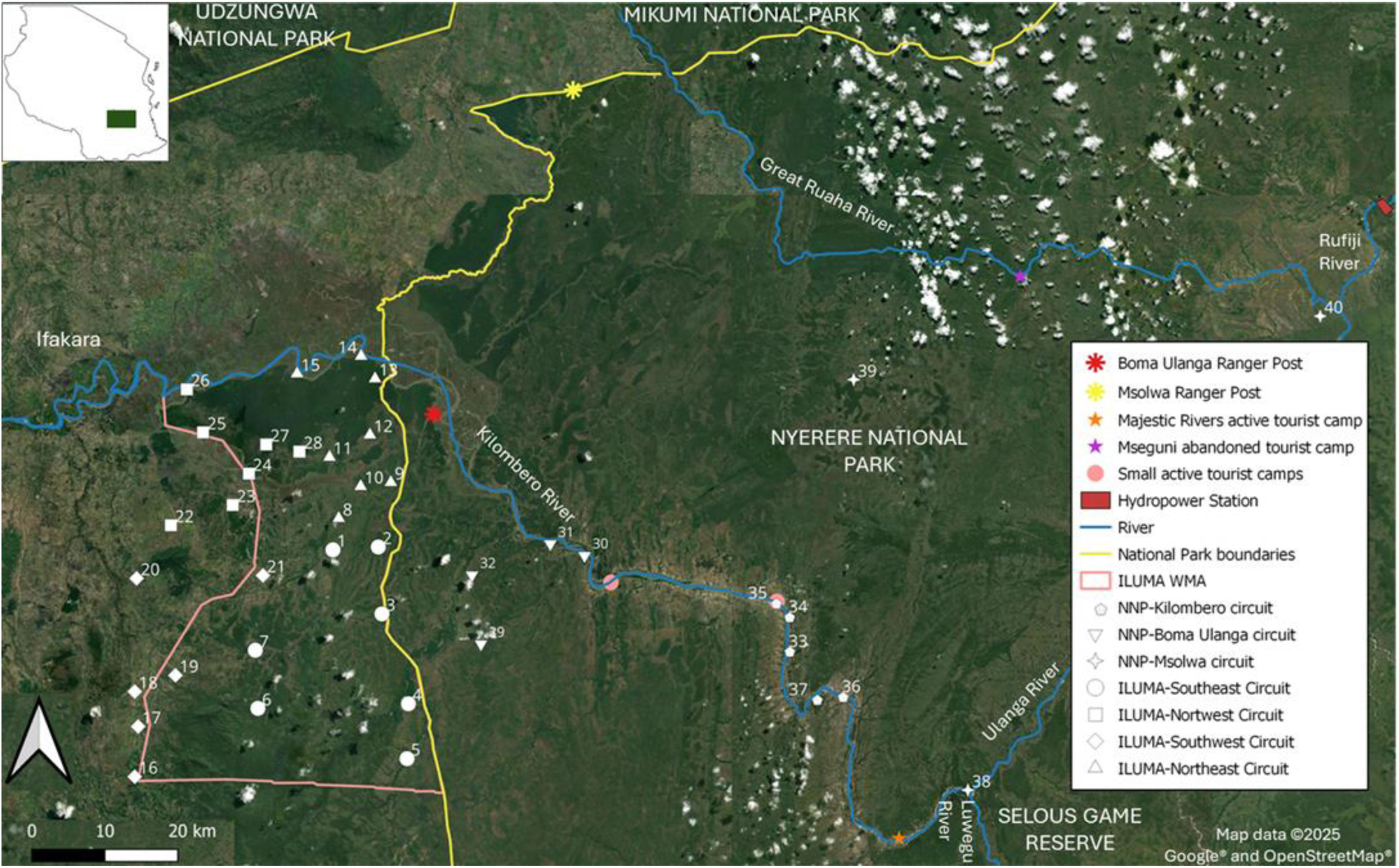
Map of the study area in its national and local context displaying the distribution of suitable camping locations used as the sampling frame for the entomological surveys of mosquitoes (Kavishe et al., 2025A; Kavishe et al., 2025B; Walsh et al., 2026) that the extension of the *VBDs360* platform described herein was developed to support the implementation. Each of the 40 camps detailed in supplementary File 1 (Walsh et al., 2026) are illustrated in the geographic context of the boundaries of the Ifakara-Lupiro-Mang’ula Wildlife Management Area (ILUMA WMA), Nyerere National Park (NNP) and the Udzungwa Mountains National Park. Map data ©2025 Google, obtained from Google Earth® and reproduced here under conditions of fair use as defined in the Google Terms of Service. Map layers were produced using base maps obtained from OpenStreetMap® under the Open Database Licence.

Live adult mosquitoes were captured with four Centres for Disease Control and Prevention (CDC) light traps (John W. Hock Company, product number 512) that were operated overnight from 19:00 to 07:00, each deployed in a distinct location in and around the camp: One suspended from a tree in an open valley or natural glade far away from humans and about 100 meters from the camp, one next to a streambed 30 to 50 meters from the camp, one at the periphery of the camp site, and one right next to a human-occupied tent.

In addition, a netting barrier interception screen was installed in a nearby open valley or natural glade, approximately 100 to 200 meters from the camp (Kavishe et al., 2025A). Collections at the interception screen were conducted using a Prokopack aspirator (Maia et al., 2011) at five time points after dusk once every hour between 19:30 and 23:30, and once before dawn at 05:30. Each collection was stored in a separate, time-labelled cup indicating the date and hour of sampling.

Surveys of mosquito larvae within water bodies at and around each camp were conducted as describe in detail elsewhere (Walsh, 2023; Walsh et al., 2026). These larval surveys were limited to within a 2 km radius of each camp location, to prevent geographic overlap between habitats surveyed from neighbouring camp locations, and to minimise potential spatial autocorrelation effects. Starting at the camp, larval surveys extended outwards until the 2km limit was reached or 4 hours has passed, starting initiated at the nearest waterbodies known to the team of Village Game Scouts, who escorted the investigators and also helped them simultaneously survey various activities of humans, livestock and wildlife (Duggan, 2023; Duggan et al., 2024; Walsh et al., 2026).

### Data recording and specimen labelling procedures for capture, transport and sorting of wild adult mosquitoes (Assay number 1)

Detailed data were recorded at the point of collection of wild adult mosquitoes in the field on a revised study-adapted standardized *assay design* (AD1) forms for each night of collection by the mobile field team. These forms were tailored to this specific study by editing standardized templates similar to those described previously (Kiware et al., 2016) include date, time, location (GPS coordinate and habitat type for the larvae), the collector’s name, and the trapping method used. For the four different ways of deploying CDC light traps and the barrier trap, each individual collection was placed in a single cup of mosquitoes, which was labelled with a primary unique key composed of the *assay design* (AD) form’s serial number (SEN), and *assay number* (AYN=1), and the *form row* number (FR) corresponding to the relevant line of data on that form. For example, a collection might be labelled AD1SEN-AYN-FR, where the actual values for these three variables recorded on the relevant form and form line were entered instead of the variable names. Because these collections of live mosquitoes in the field were not sorted *in situ* by the field team, they were also labelled with a temporary alternative secondary key at the point of collection. Specifically, the relevant date (DT) of collection and capture technique used (Coded in the forms as a combination of numerical values for the variables *Method* (ME) and *Habitat Type* (HT)) at that collection night were also written on the cup for all collections with light traps. Similarly, for the interception barrier trap collection, the same information was recorded on the collection cups, together with the time of collection to distinguish the different specific capture times coded in the HT variable. The labelled cups containing mosquitoes specimens were then placed in the mosquito carrier backpack (Kavishe et al., 2025A) and transported to the field insectary for sorting.

Each mosquito collection then underwent the process of sorting based on whether they were alive or dead (DD), visually distinguishable taxonomic classification (TX), sex and abdominal status (SAS). First, any mosquitoes found dead in the cup were separated. Second, any live mosquitoes that did not appear to belong to the *An. gambiae complex* based on their visually observable morphology (Coetzee, 2020; Gillies & Coetzee, 1987) were removed and killed. The insectary team then sorted the live and dead mosquitoes, recording their sex and abdominal status (e.g. blood-fed versus gravid, partially fed or unfed), consistent with standard entomological practice (Kiware et al., 2016). A standardized *sample sorting* (SS1) form, essentially unchanged from that previously reported (Kiware et al., 2016) was then used to record the outcome of this sorting process by listing the number of mosquitoes in each category. Each distinct individual or group of mosquitoes resulting from sorting was then assigned new unique identifiers tied to that SS1 form. Specifically, primary keys at this stage combined the SS1 form’s *assay number* (AYN=1), *Serial Number* (SEN) and the *Form Row* (FR) number for that entry (SS1: AYN-SEN-FR). An independent, fully redundant corresponding secondary key was generated using a *Sample Label Code* (SLC), *Sample Type* (ST), and *Sample Identifier Number* (SID) coded numerically as either an individual mosquito or batch of two or more mosquitoes. These labels were affixed to the microcentrifuge tubes in which sorted dead specimens were preserved (Figure 5). The use of tubes labelled with both keys ensured that if one label was damaged, written incorrectly or inaccurately recorded on the corresponding form, the sample could still be traced via the alternate key.

### Data recording and specimen labelling procedures for wild larvae (Assay number 2)

Field collections of larval specimens were similarly processed and labelled except that those field data were all entered *in situ* into an integrated *Assay Design plus Environmental and/or Sample Observation* (ADESO1) form (Figure 2). This combined form removed the need for separate linkage to separate *Assay Design* (AD) and *Environmental and/or Sample Observation* (ESO) forms (Figure 2). The ADESO1 form described herein was adapted from a similar standardised form template (Kiware et al., 2016) and used by this field project to characterise aquatic habitats and record occupancy by any *Anopheles* and by members of the *An. gambiae* complex specifically (Walsh et al., 2026). Each batch sample of live larvae pooled from a one full day of surveys around a single camp was labelled with date of collection, serial number (SEN) and form row (FR) of the ADESO1 form (Primary key), as well as the sample label code and sample type (Secondary key) as per figure 2.

### Insectary propagation and insecticide susceptibility assessments of progeny lineages derived from wild-caught specimens

In order to assess their heritable insecticide resistance traits, all live-caught adult *An. gambiae* complex mosquitoes from the field were transported in a specially designed backpack (Kavishe et al., 2025A) to a field insectary established at camp number 1 in figure 1. Upon arrival, individual surviving females were propagated in captivity through two or more (*n* ≥ 2) further generations to obtain adult *F_n_* generation progeny for assessment of their susceptibility to a pyrethroid insecticide with experimental procedures that have been standardized by the World Health Organization (WHO, 2022).

Similarly, batches of field-identified *An. gambiae* complex larvae, each pooled from different habitats surveyed over a single visit to a given camp (Walsh et al., 2026), were also brought back to the field insectary and then raised to adulthood using standard rearing procedures (Das et al., 2007). These *F_0_* adults reared in the insectary from wild-collected *F_0_* larvae were allowed to emerge and mate over a period of approximately 5 days before they were then blood-fed to obtain *F_1_* eggs, just like the live wild-caught adult females described immediately above.

All live adult *F_0_* female *An. gambiae* complex mosquitoes from each distinct larval or adult collection were kept in a single cage in the insectary and provided an opportunity to blood-fed on a cow every night for up to 3 nights. Each successfully fed female was then isolated in its own oviposition cup and then repeatedly offered additional bloodmeals, to maximize fecundity of even pre-gravid mosquitoes with underdeveloped ovarioles that may require two or more bloodmeals before they can lay eggs (Riehle et al., 2006), to establish an isofemale line of *F_n_* progeny from each wild-captured *F_0_* mother). Each *F_0_* female laid eggs (*F_1_* generation) were reared through larvae and pupae to become *F_1_* adults. These *F_1_* adults were again blood-fed on a cow as often as necessary to produce *F_2_* eggs, which were reared to *F_2_* adult mosquitoes. These *F_2_* progeny, representing grandchildren of the wild-caught *F_0_* female, were then used for insecticide susceptibility bioassays (WHO, 2022) or used to propagate further subsequent generations so that enough *F_n_* adult females could be obtained for experimental assessment of such presumably heritable phenotypes. Only *F_2_* and subsequent generations of progeny (*F_n_*, where *n* ≥2) were used for these insecticide susceptibility assays, to eliminate the risk of maternal effects upon these phenotypes and ensure sufficient numbers of progeny for satisfactorily precise quantification of this trait.

Standard WHO insecticide susceptibility tests were performed on all *F_≥2_* adult mosquitoes to assess phenotypic resistance. Mosquitoes from 20 camp locations (Figure 4) that successfully survived the field rearing process through at least two generations of progeny were used for the WHO-standardized in bioassay (WHO, 2022) of phenotypic susceptibility or resistance. The camp locations were distributed throughout the study area starting from well conserved areas inside Nyerere National Park (NNP) to the nearby villages with well-established human habitations (Figure 4). We followed the WHO tube test protocol (WHO, 2022), exposing adult mosquitoes to pyrethroid-treated papers for a fixed duration. Batches of 20 to 25 *F_n_* females were placed in standardized tubes lined with papers impregnated with a discriminating dose of a widely used pyrethroid insecticide, specifically lambda-cyhalothrin. After a one-hour exposure, mosquitoes were transferred to holding tubes with access to sugar. Knockdown was recorded at the 60-minute mark and mortality was recorded 24 hours post-exposure (WHO, 2022). Concurrently, control groups (exposed to untreated papers) were also carried out to account for any low levels of background mortality that were unrelated to the insecticide, and enable identification of experiments to be disregarded because such background mortality levels were too high (WHO, 2022).

**Figure 4.**
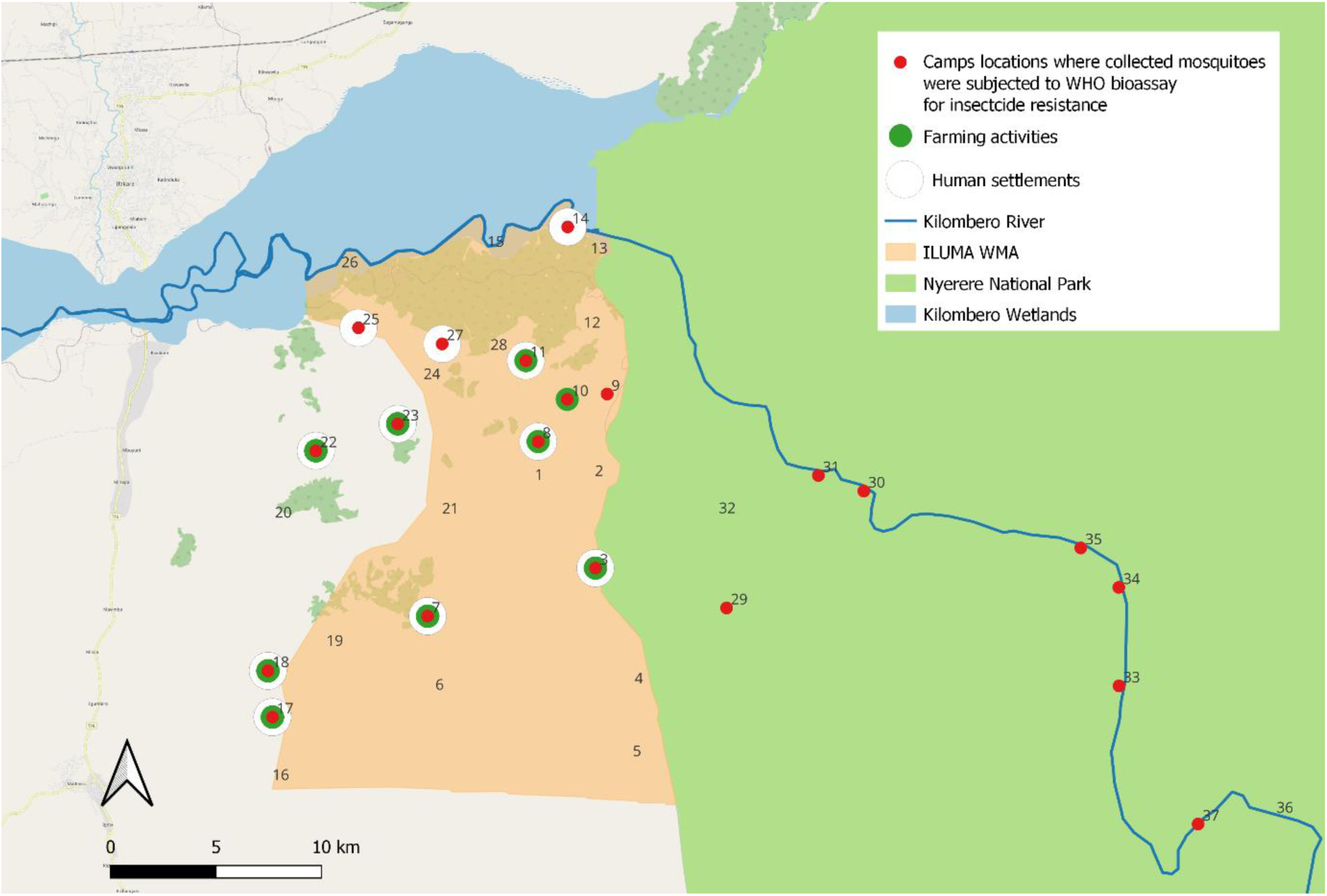
Study area map showing 20 camp locations from where *F_0_ Anopheles arabiensis* adults or larvae were collected alive, successful transported to a field insectary at camp number 1 (Kavishe et al., 2025A), and subsequently reared as independent isofemale lineages through to the F₂ or later generations. Progeny from these lineages were later subjected to standardized pyrethroid insecticide bioassays (WHO, 2022). These camp locations were deliberately chosen so that they are distributed along an ecological gradient, ranging from areas with well-conserved natural land cover, healthy wildlife populations and minimal insecticide use inside Nyerere National Park (NNP) to the east through to well-established agro-pastoral human settlements living in fully domesticated landscapes with intensive insecticide use just to the west of the Ifakara-Lupiro-Mang’ula Wildlife Management Area (ILUMA WMA).

### Data recording and specimen labelling procedures for insectary propagation and phenotype assessment of wild-caught mosquitoes (Assay number 3)

All cups containing batches of live *F_0_* female mosquitoes that survived the processes of capture, transport to the field insectary, and sorting immediately after arrival were labelled with the same primary (SEN-FR) and secondary (SLC-ST- SID) keys. These same sets of keys were then used to label all subsequently derived batches of new stages (eg. Cages of adults from trays of larvae or batches of eggs from an individual adult in a cup) and new generations (*F_1_*, *F_2_* etc) from those wild-caught *F_0_* batches.

For the standardized WHO insecticide susceptibility bioassay (AYN=3) that was then applied to *F_2_* and later generations of progeny (*F_n_*, where *n* ≥2), these same keys were used to label and distinguish individual distinct lineages of progeny that were tested. These independent, fully redundant lineage-specific identifiers were also recorded on a second set of linked assay design (AD2) and sample sorting (SS3) forms (Kiware et al., 2016). Specifically, a standardized assay design form based on the standard AD2 template was used to first record exactly what *F_n_* progeny lineage derived from what *F_0_* field-caught wild stock, or insectary colony stock, were exposed to what insecticides, when and for how long. Then a sample sorting form based on the standard SS3 template was used to record how mosquitoes subsequently self-sorted based on times to knockdown and death. Just as for the field-caught adult samples, the primary keys linking the AD2 data to the SS3 data and vice versa, respectively, were SEN-FR to SSEN-SFR (Figure 2). Similarly, each sample of mosquitoes obtained from the self-sorting process inherent to these post-exposure mortality experiments could be identified based on unique identifiers define by either the assay number, serial number and form row (AYN-SEN-FR) and or the sample label code, sample type, and sample identifier number (SLC-ST-SID). These two complementary, independent, fully redundant sets of variables respectively represented the primary and secondary keys that enabled tracing of these specimens as they proceeded through the workflow into sample storage, analysis and archiving.

### Molecular analysis and archiving of dead mosquito specimens in the laboratory

All adult mosquitoes identified as members of the *An. gambiae* complex based on standard morphological characteristics (Coetzee, 2020; Gillies & Coetzee, 1987), were stored over silica in microcentrifuge tubes. They were stored after they died during capture or were killed following the field collection and insectary procedures described above. The samples were then transported to the molecular analysis laboratory at the nearby Ifakara branch of the Ifakara Health Institute. Many mosquitoes that died during the capture process in the field or the propagation and experimentation procedures were stored as batches of several specimens placed in a single tube. However, all field-caught *F_0_* adults from which *F_n_* lineages of progeny were successful derived were stored individually in their own tube. All individually stored wild-caught *F_0_* adults, and representative samples of 5 to 10 specimens from batches of either *F_0_* ro *F_n_* adults derived from wild-caught larvae or adults, were tested for sibling species identity by polymerase chain reaction (PCR) analysis (Scott et al., 1993; Wilkins et al., 2006). All such individual or batch samples in microcentrifuge tubes were then placed inside one of 28 standard laboratory storage boxes with 81 defined positions in a 9 × 9 grid (Kiware et al., 2016). The samples were archived in order of collection, preparation and handling along two shelves in a walk-in refrigerator.

### Data recording and specimen labelling procedures for mosquito sample storage, analysis and archiving in the laboratory

All samples of mosquitoes from all of the above procedures, regardless of which distinct procedural assay they originated from either died during those collection, transport and manipulation processes, or were killed to enable their storage, analysis and archiving in the laboratory. Each sample was placed in a microcentrifuge tube and investigators then filled out the standard *Sample Observation* (SO1) form designed for recording the origins of the mosquito sample in question, where it was placed within the storage box and, later on in the laboratory, the characteristics attributed to it by molecular analyses (Kiware et al., 2016), such as polymerase chain reaction (PCR) (Scott et al., 1993), Enzyme-linked immunosorbent assay (ELISA) (Burkot et al., 1984) and blood meal identification (Burkot et al., 1981) test results.

Note that this slightly adapted version of the established SO1 form template (Kiware et al., 2016) was filled with the values for the primary and secondary key variables written on the relevant ADESO1, SS1 or SS3 form (AYN=1, 2 and 3, respectively) the sample was last associated with. When recorded accurately, each of these two alternate keys unambiguously allowed each mosquito or batch of mosquitoes to be traced back to a specific line of data on that respective source form, and these are the same variable values that were written on the labels of the microcentrifuge tubes in which the samples were preserved (Figure 5). The labeled microcentrifuge tubes containing the specimens were placed in specific positions inside the standard 9 × 9 sample storage boxes (Figure 5) that matches the specific line on the 81-line SO1 form, each of which has a serial number (SEN) that the corresponding storage box was labelled with. Conveniently, each particular form row (FR) on the SO1 form detailed exactly which row and column inside the box each sample was to be placed in and each S01 form was inserted inside the box it matched to before being transported to the laboratory for analysis and archiving.

**Figure 5.**
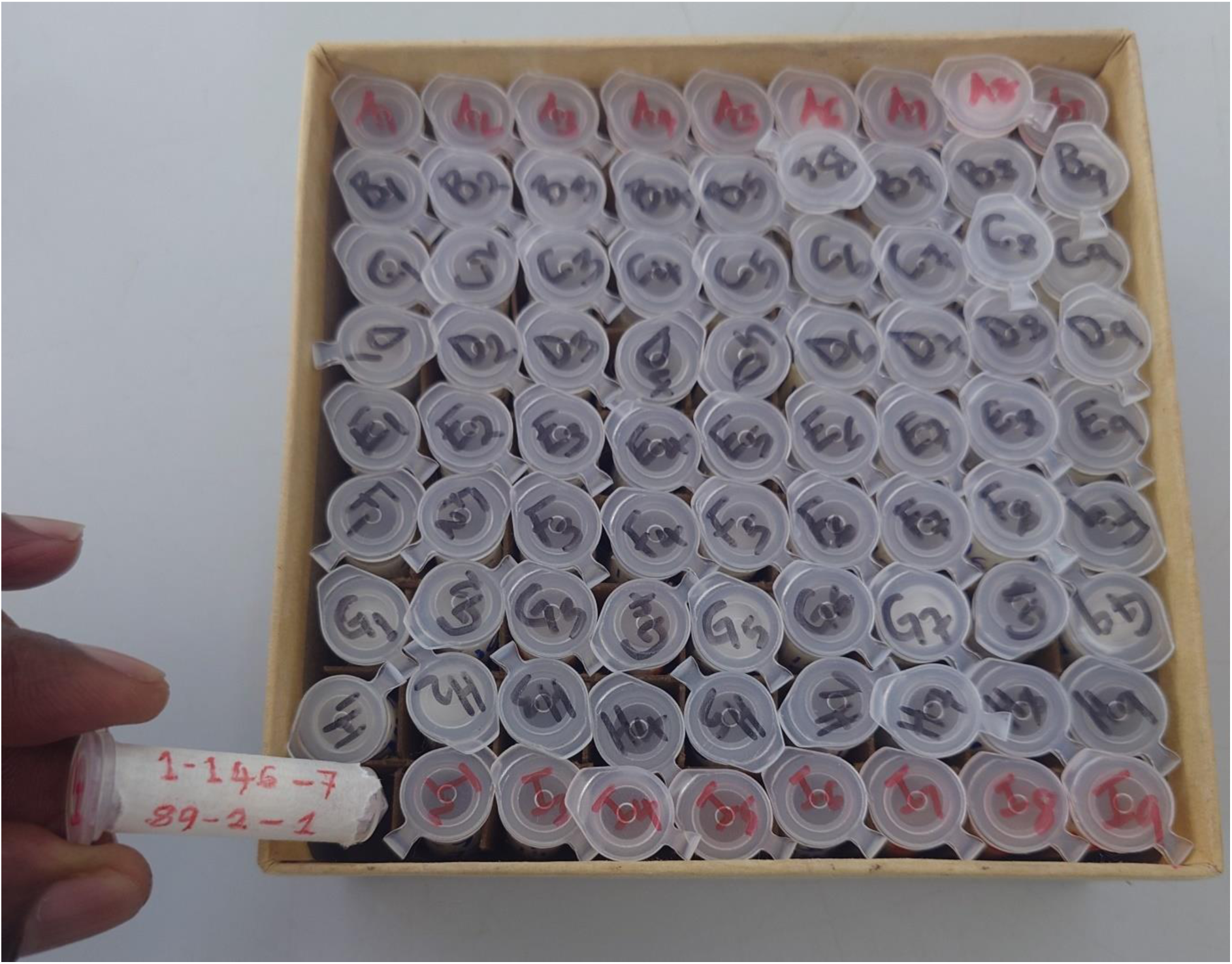
Illustration of labelling and storage workflow for dead mosquito specimens following initial field collection *Assay Design* (AD1) and *Sample Sorting* (SS1). Each microcentrifuge tube containing sorted specimens was labelled with two rows of identifiers: the first row contained assay number, serial number and form row (AYN-SEN-FR, e.g.,1-146-7) and the second row contained sample label code, sample type and sample identifier (SLC-ST-SID, e.g., 89-2-1). These identifiers were cross-referenced in the *Sample Observation* (SO1) form for subsequent laboratory analyses. Labelled tubes were organized into 9 × 9 sample storage boxes; each assigned a serial number matching its corresponding SO1 form and recorded in a Sample Storage (ST) form to ensure traceability from field data to archived specimens.

### Data analysis

The purpose of this motivating case study was to assess whether insecticide susceptibility or resistance phenotype could be related to where, when and how it was caught, with specific emphasis on geographic location, land use patterns, mammalian blood host guild composition and natural ecosystem integrity. Proportional mortality exhibited by *F_≥2_* progeny of wild-caught *F_0_* mosquitoes of unambiguously defined origin (AYN=1 or 2) in standard WHO bioassays (AYN=3) was therefore estimated for each lineage by fitting a binomial generalized linear mixed model (GLMM) without an intercept and using a logit link function, where the response was the number of dead versus surviving mosquitoes (weighted by the total tested) and the predictor was the lineage identity, included as a fixed factor. Experimental replicate units were included as random effects to account for non-independence within replicates. Model coefficients on the logit scale were back-transformed to probabilities using the inverse-logit function to obtain predicted mortality proportions (with associated standard errors) for each line.

However, given that this report specifically focuses on the fidelity of the data linkage and sample tracing processes across this complicated workflow (Figures 3 and 4), the primary outcomes are simply the proportions of such phenotype results that can be unambiguously linked back to a single specific lineage with fully defined data regarding exactly where, when and how it was collected, as well as the proportion of mosquito specimens with known mortality phenotype outcomes that can be unambiguously traced to a single mosquito specimen that can be readily found in the sample archive without manually rummaging through boxes and tubes one at a time. Secondary outcomes include what proportions of such successful data linkage and sample tracing tasks can be accomplished with the primary keys alone, without requiring the secondary keys to correct failed linkages, not only through the whole workflow but also specific pairings of tables.

## Results

The expanded informatics system provided unequivocal data connection and sample traceability at all steps of this multistage entomological experiment. A total of 22 autonomous lineages resulting from 9 wild-caught adult females and 13 larval batches were successfully reared in the field insectary to at least the *F₂*generation and exposed to standard WHO pyrethroid susceptibility bioassays. For every one of these 22 field-derived insectary lineages, absolutely all of the recorded resistance phenotype data (>8,000 *F_n_* mosquitoes tested in batches of 20 mosquitoes per test) could be clearly traced back to their exact field origin (Table 1), with fully linked data detailing where, when and how each original specimen was caught (Figure 6). The use of independent dual identifiers for each specimen or batch served as a safety net against data loss, whereby any error in the former could be corrected with the latter to recover the linkage.

**Figure 6.**
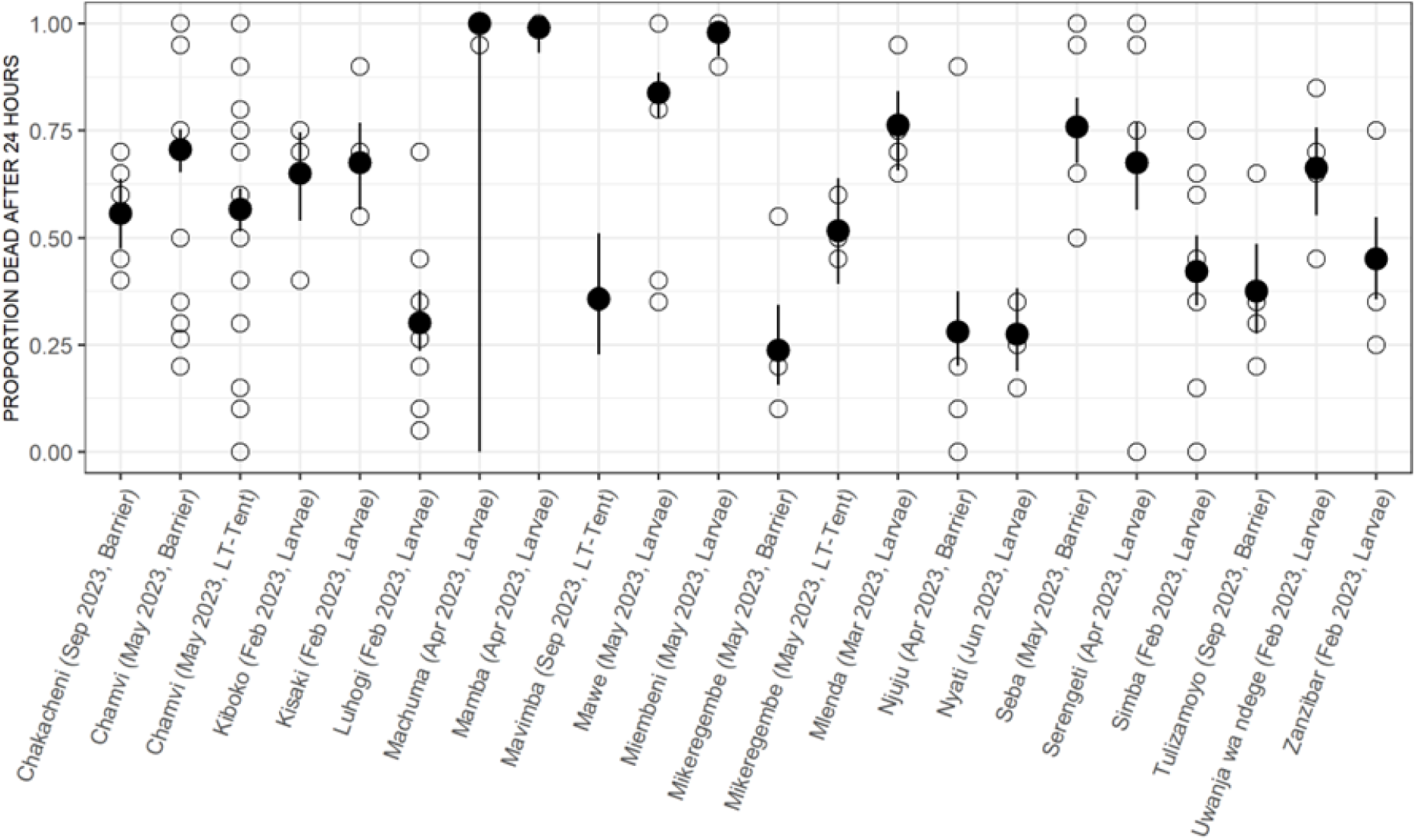
Proportional mortality outcome from the WHO insecticide bioassay of individual progeny lineages derived from distinct collections of wild mosquitoes linked to where, when, and how the original specimens were captured. The key performance outcome of the system is its ability to achieve high-fidelity data linkage and robust sample traceability of a multistage entomological experiment.

**Table 1.** Data table and primary and secondary keys for linking showing overall performance in terms of percentage linked with primary and secondary keys when used individually without cleaning and after subsequently using to cross-correct each other.

| Table | Key | Successful unambiguous linkages |
| --- | --- | --- |
| *Linkages within assay 1: Adult $F_0$ collection followed by sorting based on morphological criteria* |  |  |
| AD1 to SS1 | AYN-SEN-FR to ASY-SSEN-SFR (Primary) | 95.8% (63378/66,108) |
|  | AYN-SEN-DSEN to ASY-SSEN-SEN (Secondary) | 99.6% (65,855/66,108) |
|  | Primary cleaned with secondary and vice versa | 100% (66,108/66,108) |
| *Linkages of assay 1 to assay 3: Adult $F_0$ capture through to 9 surviving lineages of $F_n$ progeny* |  |  |
| SS1 to AD2 | AYN-SEN-FR to AYN-SSEN-SFR (Primary) | 100% (100/100) |
|  | AYN-SLC to SAYN-SSLC (Secondary) | 100% (100/100) |
|  | Primary cleaned with secondary and vice versa | 100% (100/100) |
| *Linkages of assay 2 to assay 3: Larval $F_0$ collections through to 13 surviving lineages of $F_n$ progeny* |  |  |
| ADESO1 to AD2 | AYN-SEN to SAYN-SSEN (Primary) | 100% (243/243) |
|  | SLC-ST to SSLC-SST (Secondary) | 100% (243/243) |
|  | Primary cleaned with secondary and vice versa | 100% (243/243) |
| *Linkages within assay 3: Experimental exposure of $F_n$ progeny and then sorting based on mortality responses* |  |  |
| AD2 to SS3 | ASY-SEN-FR to ASY-SSEN-SFR (Primary) | 89.7% (5070/5654) |
|  | ASY-SEN-DSEN to ASY-SSEN-SEN (Secondary) | 99.5% (5628/5654) |
|  | Primary cleaned with secondary and vice versa | 100% (5654/5654) |
| *Linkages of all field and insectary assays with laboratory workflows* |  |  |
| SS1 to S01 | ASY-SEN-FR to SASY-SSEN-SFR (Primary) | 92.1% (1826/2,022) |
|  | ASY-SLC-FR to SASY-SSLC-SFR (Secondary) | 93.7% (1895/2022) |
|  | Primary cleaned with secondary and vice versa | 97.3% (1967/2,022) |
| SS3 to S01 | ASY-SEN-FR to SASY-SSEN-SFR (Primary) | 94.0% (414/440) |
|  | ASY-SLC-FR to SASY-SSLC-SFR (Secondary) | 97.5% (429/440) |
|  | Primary cleaned with secondary and vice versa | 99.3% (437/440) |

Performance evaluation of the data linkage confirmed consistently high data fidelity across all stages of the workflow (Table 1). At the first linkage step, connecting adult field capture records to insectary sorting records (AD1 to SS1 forms), over 95% of entries matched by primary key alone and over 99% by the secondary key alone. Cross-checking between both keys resolved the few remaining discrepancies, achieving absolutely complete linkage of all >66,000 lines of data relevant to field-captured mosquitoes sorted into various taxonomic, sex and physiological status categories (Table 1).

All linkages associated with propagating the next-generation mosquitoes were also flawless after cross-correction of the independent primary and secondary keys (Table 1). Both adult-derived and larval-derived linkages of wild-caught *F_0_* mosquitoes, respectively from assay number 1 and assay number 2, to their corresponding lineages of *F_n_* progeny characterized by assay number 3 were of high fidelity before cross-cleaning and absolutely perfect afterwards (Table 1). Indeed, all data relating to each of the 9 iso-female lines derived from wild-caught *F_0_* adult females and each of the 13 lineages derived from batches of wild-collected *F_0_* larvae (13 lines) linked perfectly from the field and insectary phases (SS1 or ADESO1 forms, for assay 1 and assay 2, respectively) into the corresponding assay design records for assay 3 bioassays upon *F₂* adults (AD2 form). During the insectary bioassay phase itself (Assay 3), the link between the *F₂* assay design (insecticide exposure) records and the post-exposure outcome records (AD2 to SS3) was only 90% using the uncleaned primary key alone but exceeded 99% using the uncleaned secondary key alone, so every single one of these >5,600 records were accurately matched after cross-cleaning using both sets of identifiers to correct errors in the other (Table 1).

However, while linking the sorted field and insectary samples to the laboratory-based species identification records was also reasonably successful, but it was not entirely perfect (Table 1). Only 92% of records linking SS1 data for *F₀* specimens to SO1 records describing their sibling species identity in the molecular laboratory matched exactly via the uncleaned primary identifiers alone, and only 94% matched via the uncleaned secondary identifiers alone. After cross-correcting mismatches in the two sets of independent identifiers, >97% of the *F₀* specimens sent to the laboratory could be unambiguously linked back to their field collection data (Table 1). In other words, fewer than 3% of the *F₀* samples had any remaining ambiguity in linkage after this cross-checking. Likewise, connecting the insectary bioassay data to the laboratory analysis records for *F_n_* progeny (SS3 to SO1 linkage) was only 94% successful with the uncleaned primary identifiers alone and >97% with the uncleaned secondary identifiers, reaching >99% linkage after reconciliation (Table 1). While the few residual mismatches, occurring at the stage of transfer from the insectary to the laboratory (3 out of 440 *F_n_* samples, and 55 out of 2,022 *F₀* samples), indicate high fidelity of data integration after cross-cleaning, it is notable that these are the only imperfections documented over this entire complex combination of multiple workflows.

## Discussion

This revision and extension of the original generic schema (Kiware et al., 2016), to accommodate multiple distinct entomological assays (AYN = 1, 2, 3 etc) and include an Environmental and/or Sample Observation (ESO) table proved critical for handling more complex entomological workflows. These enhancements addressed limitations in the original schema and accommodated convenient simplifications based on study-specific idiosyncrasies, like sorting of larval collections that yields a single predefined category. This flexibility ensures that the generic schema can be adapted to study-specific adaptations while remaining both logically consistent and operationally efficient across a broader range of complex study designs. Near-perfect data provenance was preserved and transparently documented at each step, with almost all mismatches through either of the two keys resolved with the other (Table 1). The combination of linked assay design (AD) forms and sample sorting (SS) forms each with cross-referenced identifiers established a strong relational bridge between the raw field data (detailing where, when, and how each mosquito was collected) and the physical specimens that were eventually tested and stored in the laboratory. The outcome is that almost any given specimen or result can be queried to retrieve its entire history from capture and rearing background to laboratory processing with confidence in the accuracy of those linkages.

There nevertheless remains considerable room for improvement, specifically relating to the transfer of physical samples from the field insectary to the laboratory; Indeed, all of the relatively few unresolved linkage failures occurred between either assay 1 or assay 2 and assay 3. This confirms the broader experience across the Ifakara Health Institute that data linkage and sample tracing become most challenging at the interface between teams. Specifically, difficulties arises when transitioning from field and insectary teams who implement highly study specific, hands-on procedures with live mosquitoes to the laboratory team. The laboratory team typically processes larger numbers of essentially anonymized samples from lots of different studies, working in a service provide role that puts in-depth understanding of the study design out of their realistic terms of reference. While there were relatively few examples in this study of such persisting linkage failures, even a single failure to identify just one specimen of interest could badly compromise the overall success of the study if that particular *F_n_* lineage expressed an especially interesting phenotype and/or an *F_0_* specimen it was derived from proved to have an especially interesting genome sequence.

Correspondingly, an especially critical stage for maintaining traceability that received particular priority from the outset was the laboratory phase involving molecular assays and sample archiving. Correct handling of specimens, careful labelling, and strict chain-of-custody procedures are crucial in biobanking and entomological research alike, to avoid loss of data, samples or the unambiguous linkages between them (Alkhatib & Gaede, 2024). In the system reported herein, molecular processing was managed in much the same manner as biobank sample management: each mosquito tube was labelled with more than one set of identifiers, and almost every DNA extraction, PCR result, and storage location could be traced back to the relational database. Laboratory staff could retrieve almost any preserved specimen within minutes because the database recorded each mosquito’s exact tube ID and its position in the 9×9 storage box, all traceable via either the primary or secondary keys and either verifiable or correctable by combining the two. This eliminated any need for manual searching on a box-by-box and then tube-by-tube basis. This aligns with best practice in medical biobanking, where standardized processing and storage are essential prerequisites for ensuring data quality and integrity (Alkhatib & Gaede, 2024). In brief, the linkage and traceability strategy we utilized extends concepts like “Darwin Core” data standards and biobank informatics into entomology, demonstrating that insect research can achieve the same data rigor as genomics or clinical research (Wieczorek et al., 2012).

Although this practical pilot of the revised and extended generic schema for the *VBDs360* platform was implemented with paper forms to enable easy troubleshooting by the team of entomologists responsible for implementing it, it nevertheless contributes to a broader movement toward digital data management in entomology and public health. In fact, piloting the extended schema using paper-based forms was a deliberate methodological tactic to enable rigorous field testing and iterative refinement by experienced entomologists with both domain and informatics expertise. This approach allowed real-time validation of variable definitions, workflows, and data structures before digitalization, thereby minimizing design flaws and reducing downstream system redesign needs. Crucially, it also ensured that the final digital forms reflected operational realities while enhancing the robustness and usability of the extended *VBDs360* implementation. In the field of malaria vector biology and surveillance, a recent project in Ghana equipped field teams with tablet-based forms and QR-coded labels for mosquito samples, streamlining data capture and identification (Ebum, 2023). In that pilot, technicians labelled collection bags with printed stick-on labels that included both scannable QR codes and complementary alphanumeric identifiers that could be readily read by eye, and then recorded mosquito data in structured Excel® sheets on-site, enabling a subsequent bulk upload into a central database (Ebum, 2023). The PMI VectorLink Ghana team reported that this system improved sample tracking and allowed integrated data analysis across their insecticide resistance testing workflow (Ebum, 2023). Such initiatives demonstrate the feasibility of real-time, high-quality data collection even in field settings, using a combination of low-cost tools (QR labels, Excel, and an open-source DHIS2 database) (Ebum, 2023). Learning from these innovations by others, and also our own experiences with labelling and mislabelling of samples, especially under remote field conditions where printers or fully pre-printed labels may not always be available, and in laboratories where mosquito specimens may be divided into several portions that pass through different work flows, we consider that the optimal path ahead for such systems is to combine the advantages of such purely electronic data collection systems with those of the paper-based approach described herein. Specifically, while electronic data collection and management technology can be used to generate machine printable and readable primary keys that are robust within the database per se, these should be backed up with at least one simple secondary key that can be readily read by eye, manually transcribed, temporarily remembered and verbally narrated by the teams of human beings responsible for undertaking such complex studies. By leveraging mobile and cloud technologies, entomologists can drastically reduce transcription errors and delays, enabling high data quality and sample traceability that once required weeks of data cleaning in the office and manual searching for samples in the laboratory but entomological informaticians should resist the temptation to throw out the baby with the bathwater: Physical samples of mosquitoes do not reside in electronic databases that guarantee robustly perfect linkage. Instead, they pass through the hands of several fallible human beings across real-world field settings, large insectaries, and busy laboratories or archives that may house millions of specimens. Therefore, data linkages that enable sample tracing must give particular emphasis on manually applicable secondary keys. These may serve as practically robust backup to the electronic tools that are increasingly transforming entomological informatics.

Another reason to continue developing improved informatics methodologies to underpin large-scale management of entomological data and samples, is the emerging paradigm in the rise of citizen science (Južnič-Zonta et al., 2022) platforms for mosquito surveillance, which sometimes extends data handling and sample submission to unprecedented scales. The Mosquito Alert program, for instance, engages the public via a smartphone app to report mosquito sightings (Science, 2022). Users submit geo-tagged photos of mosquitoes or breeding sites, which are then verified by entomologists and compiled in a global database (Science, 2022). This effort has generated tens of thousands of occurrence records that are shared through the Global Biodiversity Information Facility (GBIF) for anyone to analyse. The integration of photos, GPS coordinates, and expert validation in near real-time exemplifies how informatics can empower large-scale, participatory science (Južnič-Zonta et al., 2022; Science, 2022).

Looking ahead, this high-fidelity data linkage can be further strengthened and generalized to enhance its applicability. A priority is real-time data collection with offline functionality, as remote fieldwork often requires paper or offline Excel records that are later uploaded to the database. Offline-capable mobile application have already been implemented within the *VBDs360* platform, allowing data to be captured in real time even in remote settings and later synchronized with central database upon availability of connectivity. Additionally, the platform supports the integration of analytical tools directly into the database, such as dashboards that calculate mortality rates or allele frequencies as data are entered, thereby transforming the system into an active analysis platform. Finally, adopting globally unique specimen identifiers, similar to GenBank accessions for sequences, would enable unambiguous cross-study sharing and reuse (Page, 2012; Page, 2008; Wieczorek et al., 2012). In this particular pilot study, the IDs were unique within our database but not globally resolvable. The biodiversity informatics community has long emphasized that stable identifiers are essential for linking records across databases (Brun et al., 2025). Piloting this concept in mosquito research, potentially through partnerships with the Global Biodiversity Information Facility (GBIF) or Barcode of Life Data System (BOLD), would ensure traceable and reusable datasets. Together, these innovations from mobile entry to universal IDs would further bridge informatics and entomology, strengthening both science and public health.

## Conclusion

In conclusion, this work present a substantial evolution of the foundational work introduced by (Kiware et al., 2016), extending its original generic schema to support more complex, multi-assay entomological experiments involving live mosquito lineages, multigenerational tracking, and cross-team workflows. This revised and extended informatics system established a rigorous chain of custody from initial field capture all the way through insectary rearing, phenotypic resistance assays, and molecular laboratory analyses. Every lineage derived from a single female adult or batch of larvae may be traced unambiguously back to records describing details of its source context and almost all of these can be traced exactly to relevant stored physical samples.

This high-fidelity linkage reduces transcription errors and preserves lineage (Wieczorek et al., 2012), allowing full historical queries of mosquito records and rapid retrieval of associated samples. Importantly, bolstering of the laboratory stage of sample handling and storage, which is often the most vulnerable point in the traceability chain, was accomplished with high fidelity through the use of dual identifiers and comprehensive recording of storage locations. As emphasized in biobanking (Alkhatib & Gaede, 2024), maintaining sample integrity and data linkage continuity in the laboratory are critical to the overall data quality. By adhering to these standards, our work shows that large-scale entomological research can meet best practices in biodiversity informatics, ensuring transparent and reproducible results across complex workflows.

While the system delivered strong performance, remaining challenges were identified at the interface between traditional project-specific entomology teams carrying out bespoke studies of live mosquitoes in the field and/or insectary and laboratory teams processing large numbers of essentially anonymized dead mosquito samples from multiple disparate projects. The way forward for addressing such persisting vulnerabilities in the chain of custody for mosquito data and samples will probably rely on the integration of such paper-based pilot systems into electronic data collection systems. This includes complementing the advantages of automatic sample labelling and mobile data entry with robust fallback procedures such as alternate (secondary) manual labelling to mitigate both practical real-world constraints upon the former and human error arising from the latter.

## AUTHOR CONTRIBUTIONS

Deogratius R. Kavishe: Conceptualization, investigation, methodology, formal analysis, writing original draft of the manuscript. Rogath V. Msoffe: Conduct and supervise bioassays, data entry and cleaning, review, editing, and validation. Selemani Mmbaga: Prepare the initial study specific schema diagrams, review, editing, and validation. Lucia J. Tarimo: Prepared the study maps, review, editing, and validation. Fidelma Butler: Review, editing, and validation. Emmanuel Kaindoa: Review, editing, and validation. Nicodem J. Govella: Review, editing, and validation. Samson S. Kiware: Revise the extended schema, review, editing, and validation. Gerry F. Killeen: Acquisition of fund, Conceptualization, investigation, visualization, methodology, formal analysis, review, editing and validation. All authors read and approved the final submitted version of the manuscript.

## ACKNOWLEDGEMENTS

The authors wish to thank the Village Game Scouts of ILUMA WMA, for their hard work and participation in field activities. We also thank all the governance, management, and stakeholder communities of the ILUMA WMA for all their collaboration and kind assistance over the course of the study. Furthermore, we thank Mr Frederic Masanja, Prof Honorati Masanja, Mr Fadhili Sango, Mr Francis Tumbo, Ms Magdalena Isaya, and Ms Elaine Kelly, Dr Ronan Hennessy, Ms Leonie O’Doherty, Ms Sonia Montero and Prof Sarah Culloty for all the essential institutional support provided by the Ifakara Health Institute and University College Cork over the course of the study. A very special word of thanks is due to our recently deceased friend and colleague, Mr Octavian Malopola, without whom this work could never have begun, much less completed safely and successfully.

## DATA AVAILABILITY STATEMENT

The data that support the findings of this study are available in the supplementary material of this article.

## ETHICAL APPROVAL

This work was conducted in accordance with all necessary ethical principles for conducting health research, and it has approvals from the Institutional Review Board of the Ifakara Health Institute (IHI/IRB/No:5-2021) and the Medical Research Coordination Committee of the National Institute for Medical Research in Tanzania (NIMR/HQ/R.8a/Vol. IX/3719).

